# Spatiotemporal control of a multilayered co-axial flow in a 3D printed microchannel with cascaded nozzles

**DOI:** 10.1101/2024.10.05.616779

**Authors:** Helen Werner, Ebrahim TaiediNejad, Mehmet Akif Sahin, Moritz Leuthner, Peer Erfle, Oliver Hayden, Andreas Dietzel, Ghulam Destgeer

## Abstract

Sculpting and stopping multilayered co-flowing streams is challenging due to inhomogeneous pressure distribution within a fluidic circuit composed of multiple interconnected microchannels having variable flow resistances. Here, we have investigated three different flow control methods to effectively stop a multilayered flow inside a 3D-printed microfluidic channel by bringing the average flow velocity from >100 mm s^-1^ to below a critical velocity of 200 µm s^-1^ within a certain delay time *t*_*D*_ of ∼2s. Firstly, we 3D printed a sequence of three concentric nozzles (∼75 µm) embedded serially inside the microchannel (∼200 µm) using a two-photon polymerization (2PP) method. Secondly, we used the 2PP-based 3D printed device to produce a structured coaxial flow of four streams with individual layer thicknesses of *O*(10 µm) within the outlet section of the microchannel. Thirdly, we removed the pressure gradient across the fluidic circuit, from > 2 bar to ∼0 bar, to stop the multilayered flow and measured *t*_*D*_ to assess the performance of the three stop flow methods. During the stop-flow phase, an inhomogeneous pressure gradient across different inlets resulted in a backflow to inlet channels with lower pressures. In the three stop-flow methods investigated, we systemically managed the fluidic capacitance to minimize a dimensionless backflow index (*BFI*) value from ∼0.3 (worst case) to ∼0.03 (best case) for a total flow rate ranging from 16.8 µl min^-1^ to 168 µl min^-1^. Finally, we have recommended the best stop-flow conditions, which resulted in a minimal delay time of *t*_*D*_ ∼ 2s and a BFI < 0.05.

## 1 Introduction

The ability to effectively control, i.e., sculpt and stop, a microfluidic flow is crucial to the fabrication of structured microparticles^[1]^ and fibers^[2]^, which have found numerous applications across the fields of cell biology^[3]^, diagnostics^[4–8]^ drug delivery^[9,10]^, micro-robotics^[11–13]^, and tissue engineering^[14]^. Planer 2.5D microfluidic devices utilizing hydrodynamic flow focusing^[15–17]^, and inertial flow sculpting^[18–23]^ have been used to shape the flow profile, but there is a limited choice for microchannel designs due to dependence on the soft lithography process. Compared to 2.5D microfluidic devices, 3D-printed microchannels offer a great diversity of microchannel designs and even greater control over the sculpted flow profiles^[24–28]^ We have previously demonstrated 3D-printed intertwined networks of microfluidic channels with an outlet hydraulic diameter (*D*_*h*_) of 400-1000 µm to sculpt multi-layered flow streams with features of *O*(100 µm)^[25–27]^. However, it was challenging to further reduce the microchannel diameter for enhanced flow sculpting capabilities due to the limitations in the 3D printing resolution. Stereolithography-based 3D printers used to fabricate the 3D microfluidic channels have a printing resolution of *O*(20 µm), which struggles to successfully fabricate channels with < 100 µm gaps and wall thicknesses. Therefore, high-resolution 3D printing of smaller channels is needed to realize sculpted flow profiles with feature *O*(10 µm).

Moreover, stopping the microfluidic flow of curable precursors is essential to manufacturing 3D particles with enhanced spatial resolution using a stop-flow lithography process^[29,30]^. The fluidic capacitance of a microfluidic system hinders an immediate stopping of the flow after the removal of the pressure gradient across the fluidic circuit^[31–34]^. We have recently investigated the effect of fluidic capacitance on the stopping of a microfluidic flow inside microchannels with *D*_*h*_ of 100-500 µm^[35]^. It was particularly challenging to stop a flow driven by a syringe pump through a single inlet microchannel with a *D*_*h*_ of 100 µm due to a relatively slower dissipation of fluidic capacitance and higher flow resistance *O*(10^14^ Pa.s/m^3^). In contrast, we could stop a multi-layered co-axial flow within a microfluidic device with four inlets merging into a single outlet channel with a *D*_*h*_ of 500-1000 µm and flow resistance *O*(10^10^-10^11^ Pa.s/m^3^)^[27]^. However, stopping a multi-layered flow within smaller microfluidic channels with *D*_*h*_ of ≤ 200 µm and higher flow resistances *O*(10^13^ Pa.s/m^3^) poses significant challenges that make it pertinent to develop new stop-flow methods.

In this work, we present a miniaturized microfluidic device with a cascaded nozzle sequence to sculpt a multi-layered co-axial flow, which was controlled by three stop-flow methods to realize the best experimental conditions (**Figure 1**). We incorporated a high-resolution two-photon polymerization (2PP) based 3D printing^[36]^ to fabricate an intertwined network of 3D microfluidic channels (*D*_*h,c*_ ∼ 200 µm) with integrated nozzles (*D*_*h,n*_ ∼ 75 µm) (Figure 1a). Three free-standing nozzles, supported by crossbar structures, were aligned concentrically within the main channel. Four independent flow streams originating from the four inlets (I_1_-I_4_) were sculpted into a concentric multilayered co-axial flow profile within the outlet cross-section aa’ of the device (Figure 1b). The thickness of individual layers was precisely tuned with a spatial resolution of *O*(10 µm) by varying the inlet flow rates (*Q*_*1*_-*Q*_*4*_). We investigated three different flow control methods, distinguished based on the mechanism of producing or removing the pressure gradient within the fluid circuit, to start or stop the sculpted flow in a cyclic manner. Firstly, we rapidly remove the pressure gradient (i.e. Δ*P =* (*P*_*in*_ - *P*_*out*_) → 0) to bring the average flow velocity (*v*_*avg*_) below a critical value (i.e., *v*_*avg*_ < *v*_*crit*_) during the stop phase of the cycle within a delay time of *t*_*d*_. Secondly, we quickly generate the pressure gradient again to stabilize the flow back to its earlier state during the flush phase of the cycle within a stabilization time of *t*_*s*_, and repeat (Figure 1c). An inhomogeneous pressure drop from inlets I_1_-I_4_ to outlet O, since *P*_*1*_ > *P*_*2*_ > *P*_*3*_ > *P*_*4*_ due to mismatched flow resistances, can lead to an undesirable residual backflow between the channels as the pressure within the channels is balanced, such as *P*_*1*_ ≈ *P*_*2*_ ≈ *P*_*3*_ ≈ *P*_*4*_ ≈ *P*_*out*_ (Figure 1d). We have investigated the degree of backflow for three stop-flow methods to identify the best experimental conditions for rapid stop-flow with minimal backflow.

**Figure 1:**
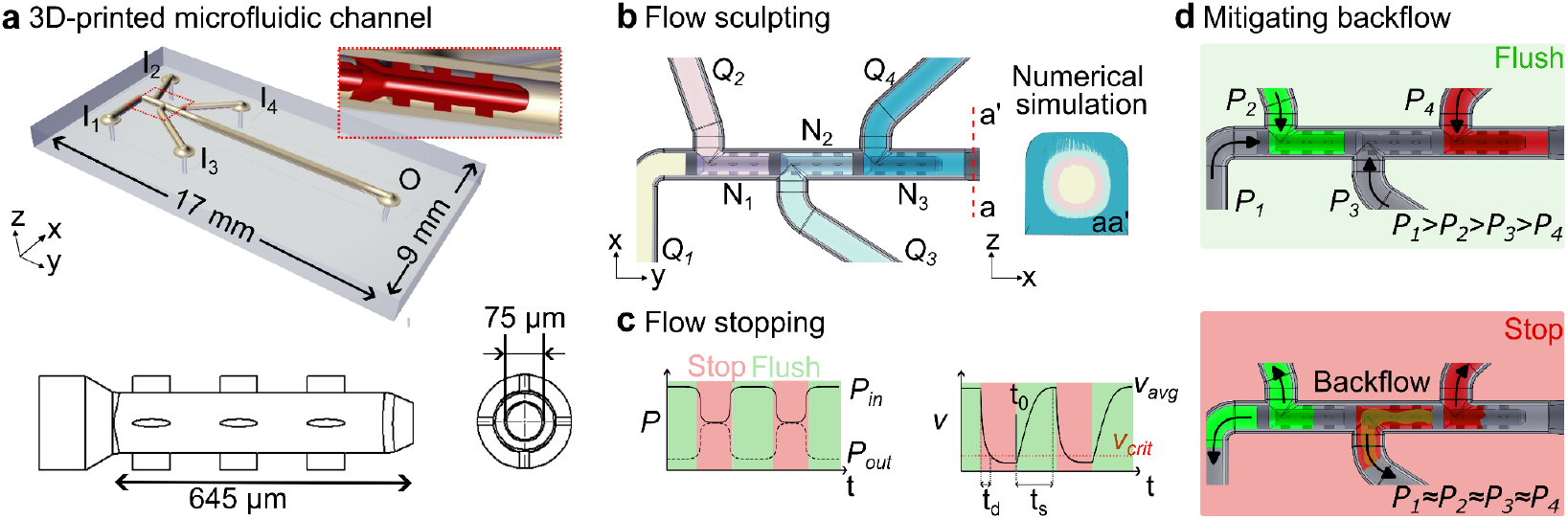
(a) Microfluidic device design with four inlets (I_1_-I_4_) and integrated nozzles. The inset shows a side view (yz-plane) of a nozzle. The drawing at the bottom depicts the top and front views of a nozzle. (b) Sculpting of four streams, pumped at flow rates *Q*_*1*_*-Q*_*4*_ through the three nozzles (N_1_-N_3_), into a concentric flow profile predicted by a numerical simulation within the outlet cross-section aa’. (c) Stopping the sculpted flow by matching the inlet and outlet pressures (*P*_*in*_ ≈ *P*_*out*_) to bring the average flow velocity (*v*_*avg*_) below a critical value (*v*_*crit*_). (d) An inhomogeneous pressure drop for the four inlets during the flush phase leads to a backflow within channels during the stop phase as the system pressure homogenizes. The arrows indicate the flow direction.

## 2 Results and discussion

### 2.1 Manufacturing of microfluidic devices with integrated nozzles

We 3D-printed a microfluidic device on a borosilicate glass substrate with channels precisely aligned with five through holes for fluidic connections, i.e., four inlets (I_1_-I_4_) and one outlet (O) port (**Figure 2**a). The 3D printed channels were covered with glue to sustain an internal high fluidic pressure and protect against external physical damage. A close-up scanning electron microscopy (SEM) image of the 3D printed channels, with the outer channel wall removed for visualization, reveals a perfectly aligned cascade of three nozzles in the middle (Figure 2b-c). To stabilize the nozzles during 3D printing and realize a concentric co-axial fluid flow, we designed our microfluidic devices with two different support structures for nozzles (Figure 2d). Preliminary microfluidic devices printed without a supportive structure around the nozzle resulted in an eccentric flow development because of the nozzle instabilities during the 2PP- based printing and bending under the inhomogeneous fluid pressure during the experiments (Figure S1a). In a revised design, we included a four-step support structure to prevent nozzle bending and to uniformly distribute the flow around the nozzles; however, the added supports resulted in an undesirable high flow resistance and pressure drop (up to ∼0.3bar) across each nozzle (Figure S1b and Table S1). Therefore, a single cross-bar-shaped support was used instead to reduce the fluidic resistance and pressure drop (by 3x to ∼0.1bar) (Figure S2 and Table S2). However, a single cross-bar was not enough to prevent the sagging of the stand- alone nozzles during 3D printing. The 2PP-based 3D printing is executed in a sequential manner to print blocks of ∼(200 µm)^3^ in order, where the absence of bottom support at the stitching planes resulted in sagging of the nozzles. Hence, we provided three supports at equal intervals of ∼200 µm or a single long support at the bottom with segregated supports at the top and sides to prevent sagging of the 3D printed nozzles (Figure 2d). Lastly, the 3D-printed device was mounted on a compatible aluminum holder, where the fluidic ports of the glass substrate and the holder were perfectly aligned (Figure 2e). Five O-rings were sandwiched between the holder base and the glass substrate by screwing an aluminum lid on top for leakage-free fluidic connections during the flow sculpting and stop-flow experiments (sections 2.2-2.4). The aluminum base has four inlet-female ports and one outlet-female port suitable for screwing male connectors for the fluidic supply and waste collection, respectively.

**Figure 2:**
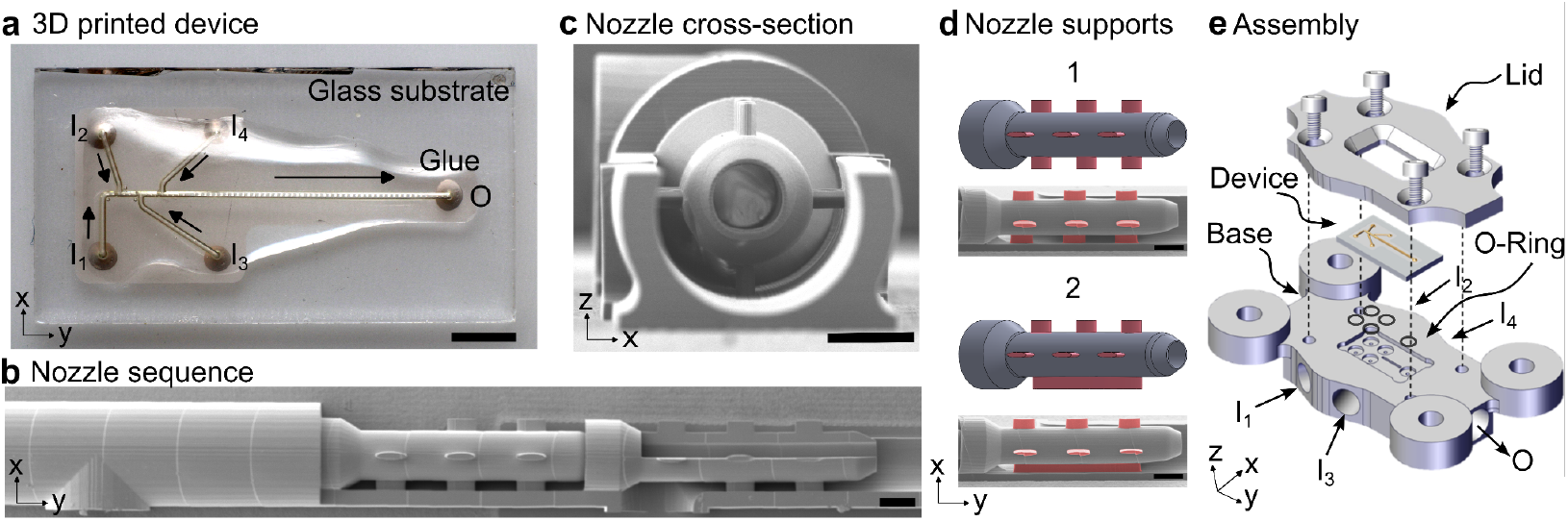
Microfluidic device 2PP printed on a glass substrate with a compatible holding system for fluidic connection. (a) Microfluidic device with four inlets and one outlet printed on a glass substrate with five holes for fluidic access. Scale bar: 2mm. The 2PP structures are mechanically stabilized by embedding them in glue. (b) SEM images of the interior nozzles within the gradually cut outer channel outer wall and nozzle. Scale bar: 100µm. (c) An isometric view highlighting the nozzle cross-section. (d) Two different types of support structures stabilize the nozzle during the printing process and avoid stitching issues during the fabrication process. Scale bars: 100µm. (e) Assembly of microfluidic device and compatible holding system.

### 2.2 Sculpting a multilayered coaxial flow

The 3D microfluidic device routes four fluid streams through a sequence of three nozzles to obtain a co-axial multilayered flow profile (**Figure 3**a). A flow stream, entering the main channel via inlet 1, is compressed through nozzle 1 (N_1_), as another flow stream from inlet 2 engulfs N_1_ from outside. Both streams meet at the tip of N_1_ and merge to form a bilayer co-flow within the bb’ cross-section with minimal diffusive mixing due to high Peclet number (Pe) *viz. O*(10^4^-10^5^). The third and fourth streams join the bi-layered flow after N_2_ and N_3_, respectively, to form concentric tri- and quadri-layered cross-sectional flow profiles, respectively (see cc’ and dd’). The flow streams 2 and 3 are colored red and green to visualize and characterize the concentric flow profile numerically and experimentally.

**Figure 3:**
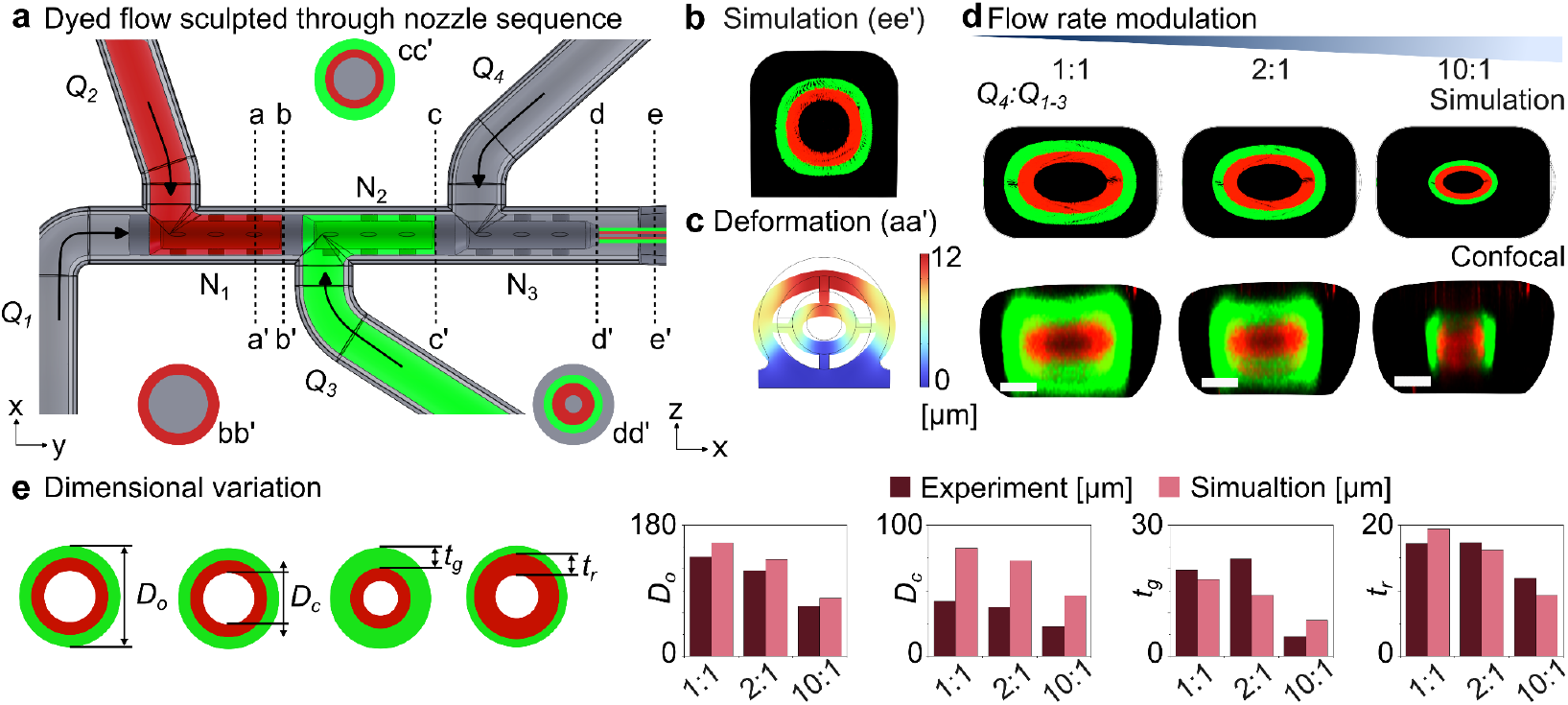
Microfluidic flow sculpting using concentric nozzles and variable flow rate ratios. (a) Top view of the microfluidic device with four inlets to introduce samples at variable flow rates (*Q*_*1-4*_) that merge into a multilayered co-flow after passing through the nozzle sequence (N_1_-N_3_). (b) Numerical simulation depicting concentric flow profile within cross-section ee’. (c) Numerical simulation showing channel deformation at cross-section aa’. (d) Modulated of cross-sectional flow profile by increasing the flow rate ratio (*Q*_*1*_:*Q*_*2-4*_) from 1:1 to 10:1. The confocal images were obtained with a total flow rate of 168 ul min^-1^ and a Reynolds number of 0.31 to ensure a perfectly laminar flow. A Peclet number of 1.8×10^5^ limited the diffusive mixing of laminar streams. Scale bars: 50µm. (e) Dimensional characterization of sculpted flow profile under variable flow rate ratios.

Numerical simulations predicted a multilayered concentric co-axial flow profile within the outlet channel cross-section ee’ after the flow sculpting by concentric nozzles (Figure 3b). In the experiment, the 3D-printed channel cross-section deformed under a load of protective glue applied on top of the channel (Figure 3c and S3), which in turn resulted in an elliptical cross-sectional flow profile (Figure 3d). The glue layer imposed external pressure on the channel walls, causing deformation during the curing of the glue. The nozzle, connected to the outer channel walls by a support structure, also deformed under this pressure. The deformation resulted in the oval-shaped nozzle tip and channel cross-section (Figure S3), which led to an elliptical flow profile confirmed by numerical simulations and confocal microscopy. Experimentally obtained confocal microscopy images for the cross-sectional flow profile matched reasonably well with the numerical simulations (Figure 3d). The flow rate ratio (*Q*_*4*_:*Q*_*1- 3*_) was increased from 1:1 to 10:1 to sculpt the flow profile and tune the individual layer thicknesses. The outermost flow rate was increased to compress the inner three streams towards the channel center, thereby, squeezing the thicknesses of these layers. The experimentally obtained outer diameter of the green layer (*D*_*o*_), inner cavity diameter of the red layer (*D*_*c*_), and thicknesses of green (*t*_*g*_) and red (*t*_*r*_) layers matched well with the numerical simulation results for variable flow rate ratios (Figure 3e; see also Table S3-S5). A decreasing trend with an increasing flow rate ratio was evident for all the dimensions. The experimental and numerical values of *D*_*o*_, *t*_*g*_, and *t*_*r*_ were reasonably close. However, the experimental *D*_*c*_ values were much lower than those obtained from the simulation, which was attributed to the diffusion of red dye towards the channel center. Notably, for a flow rate ratio *Q*_*4*_:*Q*_*1-3*_ of 10:1 inside a 200 µm outlet microchannel, we could experimentally obtain individual layer thicknesses (*t*_*g*_ and *t*_*r*_) within the sculpted flow profile with dimensions *O*(10µm). We will need a flow rate ratio *Q*_*4*_:*Q*_*1-3*_ of up to 200:1 to produce similar layer thicknesses within a 500 µm outlet microchannel (Figure S4), which would be challenging to realize experimentally.

### 2.3 Stopping a multilayered coaxial flow

A multilayered coaxial flow developed within the 3D-printed microfluidic device as the four inlets were pressurized using syringe pumps (**Figure 4**a). To rapidly stop the flow, the pressure gradient across the fluidic circuit was removed quickly by turning the syringe pumps off and switching the valves upstream and downstream of the microchannel (**Movie S1**). The fluidic capacitance of the microfluidic circuit, due to the compliance of the channel walls and the connecting tubing, hinders an immediate stopping of the flow. We used glass syringes, PTFE tubings, and a microfluidic device printed with a high Young’s modulus material to minimize fluidic capacitance. As the pressure gradient vanished, i.e., (P_in_ - P_out_) → 0, the average flow velocity (*v*_*avg*_) reached below a critical value (*v*_*crit*_ ≈ 200 µm s^-1^) within a certain delay time (*t*_*d*_), and the flow was considered stopped (Figure 4b). The critical flow velocity *v*_*crit*_ was defined as a distance traveled equal to the channel diameter per second. We measured the average flow velocity (*v*_*avg*_) by tracing fluorescent microparticles within the microchannel for three consecutive stop-flow cycles. An in-house built algorithm detected the particle positions between two subsequent frames 40 ms apart to extract the individual particle velocities, which were averaged to obtain the flow velocity.

**Figure 4:**
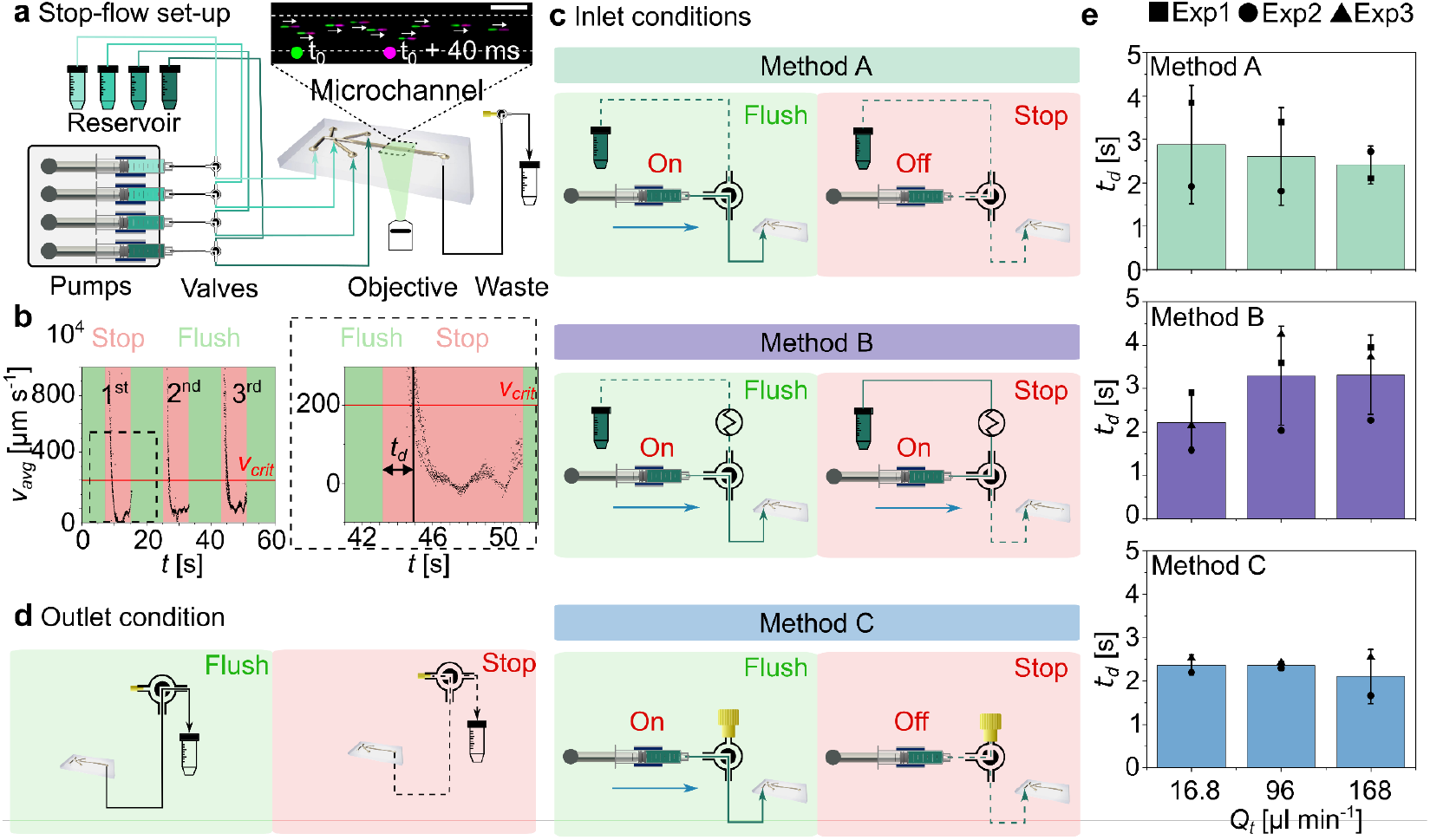
Three stop-flow configurations for cyclic flow control within a multiple inlet device. (a) Experimental stop-flow set-up including four syringe pumps connected to the microfluidic device and separate reservoirs *via* three-way valves. The outlet flow passes through another valve to a collection tube. Suspended microbeads are traced to determine the average flow velocity (inset). Scale bar: 200 µm. (b) Three representative stop-flow cycles, including stop and flush phases, depict the cyclic *v*_*avg*_ trend over time. The flow is considered stopped if *v*_*avg*_<*v*_*crit*_ after a certain delay time (*t*_*D*_). (c) Three different inlet conditions result in Method A-C. (d) Outlet condition with a two-way valve, modified from a three-way valve, is used with all inlet conditions. (e) Experimentally determined *t*_*D*_ using three different flow rates *Q*_*t*_ for Methods A-C.

We have investigated three stop-flow configurations, *viz*. Methods A-C, with different inlet conditions to characterize the delay time *t*_*d*_ during the stop phase for variable total flow rates (16.8 – 168 µl min^-1^) during the flush phase (Figure 4c-e). In front of the four syringes and at the outlet of the microfluidic device were 3-way valves directing the flow in the desired directions. At the inlet, a 3-way valve connected the syringe to the microfluidic device during the flush phase and to a reservoir or to a closed end of the valve during the stop phase (Figure 4c). For Method A, the syringes were connected to the microfluidic device during the flush phase as the pumps were running. During the stop phase, the syringe pumps were turned off as the 3-way valves linked the syringes to their respective reservoirs. This configuration enabled us to refill the syringes between experiments without removing them from the syringe pumps. For Method B, we added additional flow resistances between the valves and the reservoirs to keep the syringes pressurized during the stop phase, as the syringe pumps were kept running over the whole cycle. For Method C, one port of the 3-way valves was plugged to stop the flow during the stop phase, whereas the syringe pumps were also turned off. Similarly, a 3-way valve with one port plugged was used at the outlet for Methods A-C (Figure 4d). All three methods aimed to remove the pressure gradient within the fluidic circuit as quickly as possible to realize minimal delay times.

We measured an average delay time *t*_*d,avg*_ of 2.63 s, 2.94 s, and 2.23 s for Methods A, B, and C, respectively, for variable flow rates (Figure 4e). In Method A, since the syringes were connected to the reservoirs at the atmospheric pressure (*P*_*atm*_) during the stop phase, they were depressurized from the maximum inlet pressures of *P*_*in*_ to *P*_*atm*_. Therefore, when the inlet valves connected the syringes back to the microfluidic device, the inlet pressures had to first build up from *P*_*atm*_ to a maximum level of *P*_*in*_ before generating a stable velocity within the microchannel outlet during the flush phase (see Figure S5). For example, to generate a total flow rate of 160 µl min^-1^, the syringes connected to inlets 1-4 needed to reach a *P*_*in*_ of 1.1, 1.0, 0.7, and 0.3 bar, respectively, from *P*_*atm*_ = 0 bar. An inhomogeneous pressure drop between the inlets also caused an undesirable backflow toward the inlet channels (Section 2.4). In Method B, we increased the flow resistances towards the reservoirs to match them with the flow resistance of the microfluidic device. The syringe pumps were kept running to preserve the pressure of the fluidic circuit as the inlet valves were switched between the microchannel and the reservoirs during the flush phase and the stop phase, respectively. Since the inlet pressures were not dropping to *P*_*atm*_, this configuration enabled the flow to stabilize relatively quickly during the flush phase (see Figure S6), and the backflow (Section 2.4) was significantly reduced when compared to Method A; however, the average delay time (*t*_*d,avg*_ = 2.94 s) did not improve much during the stop phase. In Method C, we disconnected the reservoirs from the fluidic circuit to reduce the unnecessary pressure fluctuations. The syringe pumps were turned off during the stop phase, but the altered inlet valve configuration kept the syringes and the fluidic circuit pressurized. The average delay time (*t*_*d,avg*_ = 2.23 s) improved, the flow stabilized quickly during the flush phase (Figure S7), and the backflow (Section 2.4) was significantly reduced in Method C. We conclude that the high flow resistance of the microfluidic channels *O*(10^13^ Pa·s m^-3^) was the main contributor to the stop flow delay time (*t*_*d*,_), whereas, the inlet tubing and valves configurations (Methods A-C) were responsible for driving the backflow and delayed flow stabilization.

### 2.4 Backflow in a multichannel microfluidic device

An inhomogeneous pressure drop (Δ*P*_*i*_) between the inlets led to a backflow phenomenon (**Figure 5**a-b). A chain of causation can be traced to an inhomogeneous hydraulic flow resistance within the fluidic circuit. For example, the flow from inlet I_1_ sequentially passed through nozzles N_1_ to N_3_ (*D*_h,n_ = 75 µm) to reach the outlet by overcoming a relatively higher flow resistance *R*_*1*_ of 2.0×10^14^ Pa·s m^-3^, which was ∼5x higher than *R*_*4*_ of 0.4×10^14^ Pa·s m^-3^. Therefore, the inlet I_1_ was pressurized at a much higher level (1.42 bar) compared to inlet I_4_ (0.29 bar) to reach a stable co-flow during the flush phase. We performed numerical simulations to estimate the pressure drops Δ*P*_*1-4*_ between the four inlets and the outlet during the flush phase, where the highest pressure drop Δ*P*_*1*_ ≈ 5^*^Δ*P*_*4*_ was recorded for inlet I_1_. As the pressure across the fluidic circuit was homogenized during the stop phase, the backflow ensued from the high-pressure to the low-pressure segments of the fluidic circuit.

**Figure 5:**
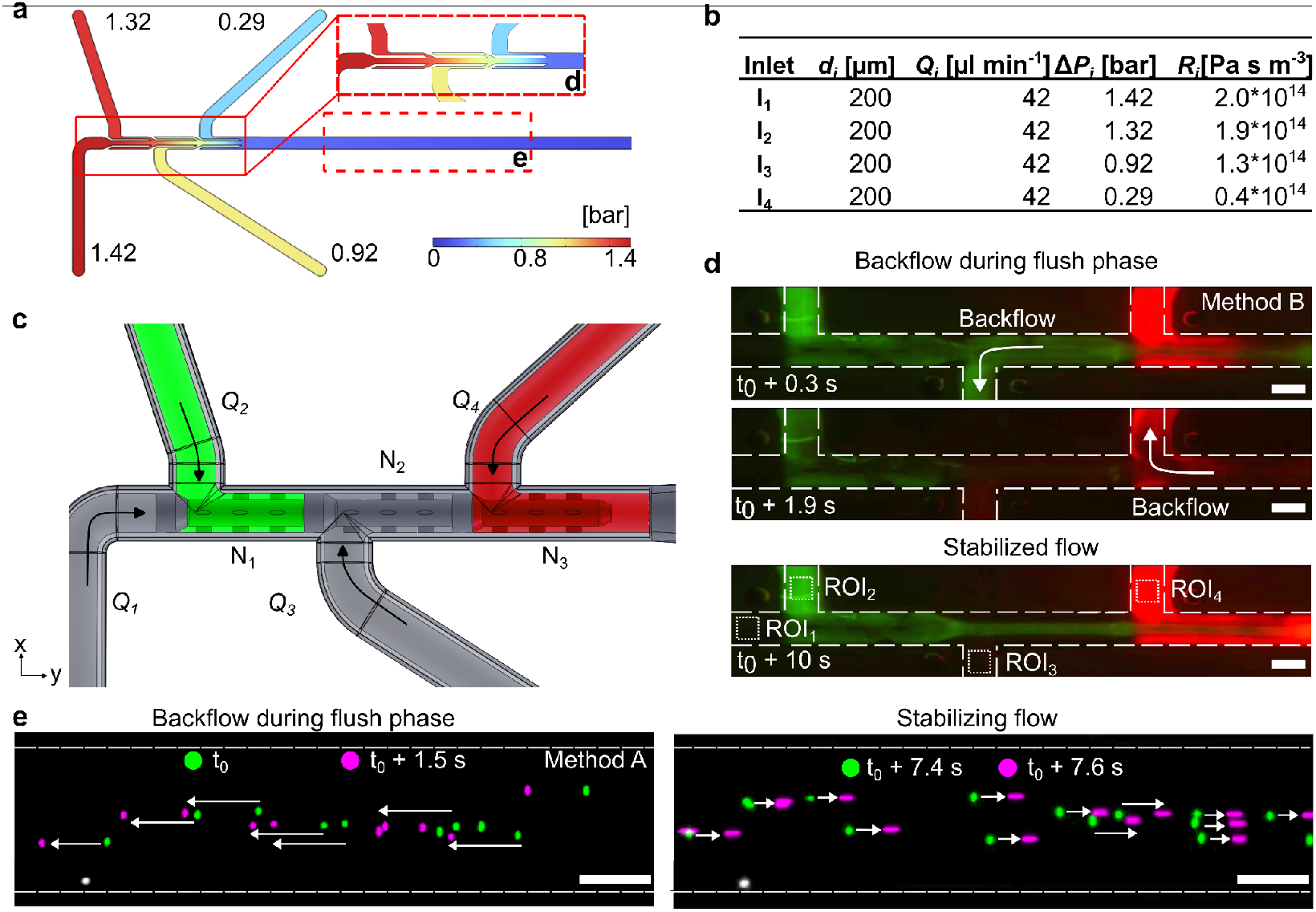
Backflow during the stop-flow cycle. (a) The numerical simulation depicts an inhomogeneous pressure drop between the inlets and the outlet of the device. (b) Simulation parameters and resulting pressure drop for each stream. (c) We use green (*Q*_*2*_) and red (*Q*_*4*_) color streams together with the transparent (*Q*_*1*_ and *Q*_*3*_) streams to trace the backflow. (d) For example, using Method B with *Q*_*t*_ of 16.8 µl/min and flow rate ratio *Q*_*4*_:*Q*_*1-3*_ of 1:1, the backflow is evident at up to 1.9 s into the flush phase. A stabilized flow is recorded at 10 seconds into the flush phase. Four regions of interest (ROI) are defined for fluorescent intensity measurement for subsequent analysis. (e) The overlayed images show fluorescent tracing beads flowing with the backflow early on during the flush phase of Method A and moving forward as the flow starts to stabilize later on. The green and purple colors indicate the time difference between the two frames, and the white arrows indicate the velocity magnitude. The scale bars are 100 µm.

For a quantitative analysis of the backflow, we performed stop-flow experiments by introducing green and red fluorescent dyes in flow streams 2 and 4, respectively (Figure 5c and **Movie S2**). The flow was going back into non-specific inlets during the flush phase when it was supposed to move forward toward the outlet. For example, it was evident during the stop-flow Method B operated at a relatively low flow rate of 16.8 µl/min that the flow was going back to the inlets after several seconds into the flush phase, and it took up to 10 seconds for the flow to fully stabilize (Figure 5d). The flow stabilization was relatively faster at higher flow rates. We recorded the fluorescent intensity at four regions of interest (ROI_1-4_) within the microchannel to quantitatively analyze the backflow phenomenon. Moreover, particle tracing within the main channel of the device also confirmed the backflow during the flush phase of Method A despite the pumps running in the forward direction (Figure 5e). The tracing beads only moved in the forward direction after >7s the pumps were turned on for the flush phase.

The average flow velocity (*v*_*avg*_), obtained using the particle tracing, decreased below the critical velocity (*v*_*crit*_) after a certain delay time during the stop phase; however, a negative flow velocity and a delayed response towards flow stabilization were observed during the flush phase, which was a clear indication of backflow within the microfluidic device (Figure 6a). For a quantitative analysis of such backflow, we measured the fluorescent intensities for the green and red dyes mixed in two of the four co-flowing streams and separately analyzed them for Method (A-C) for multiple flow rates to obtain the *BFI* (Figure b-e). The *BFI* was defined as the sum of *BFI*_*G*_ and *BFI*_*R*_ associated with the green and red channels of the captured RBG images of ROI_1-4_, respectively (equation 1).

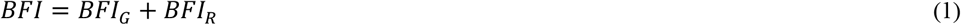

**Figure 6:**
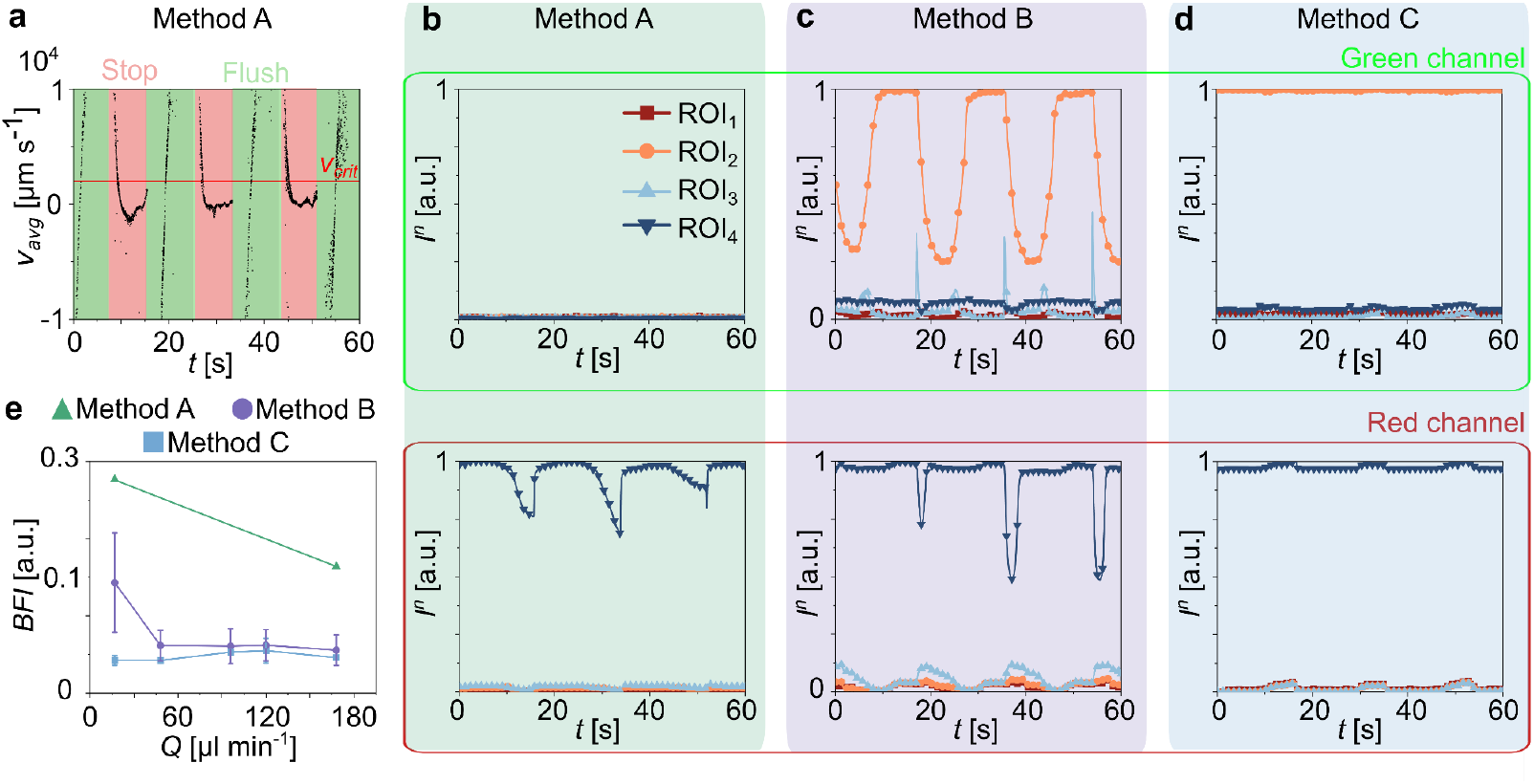
Quantitative analysis of backflow during the stop-flow cycle. (a) Average flow velocity over three stop- flow cycles, using Method A with *Q*_*t*_ of 16.8 µl min^-1^ and *Q*_*1*_:*Q*_*2-4*_ = 1:1, shows a negative flow velocity (i.e., backflow) during the flush phase. (b-d) Normalized green and red fluorescence intensities (*I*^*n*^) were measured for ROI_1-4_ for Method A-C with *Q*_*t*_ of 16.8 µl min^-1^ and *Q*_*1*_:*Q*_*2-4*_ = 1:1. (e) Backflow index (BFI) for different flow rates for Method A-C.

The *BFI*_*G*_ was a sum of the areas under the curves for the normalized green fluorescent intensities plots, i.e., *I*^*n*^(ROI_1,3,4_) and the area over the cure for the normalized green fluorescent intensity plot *I*^*n*^(ROI_2_) over a maximum time *t*_*max*_ (Figure 6b-d). The sum was normalized by 4 x maximum normalized fluorescent intensity (*I*_*max*_ = 1) and the maximum imaging time (*t*_*max*_ = 60s).

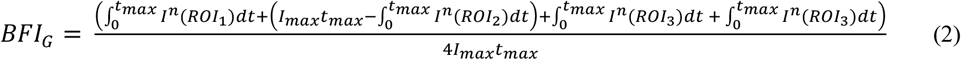

The *BFI*_*R*_ was also a normalized sum of the areas under the curves for *I*_*ROI_1,2,3*_ and the area over the cure for *I*_*ROI_4*_ over a maximum time *t*_*max*_ (Figure 6b-d).

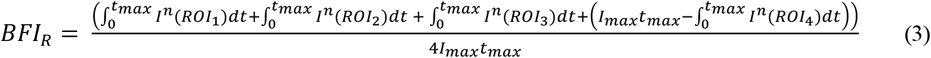

In an ideal case without any backflow, i.e., *BFI* = 0, we shall obtain *I*^*n*^(ROI_2_) = 1 and *I*^*n*^(ROI_1,3,4_) = 0 in the green channel, and *I*^*n*^(ROI_4_) = 1 and *I*^*n*^(ROI_1,2,3_) = 0 in the red channel. For a worst-case scenario, i.e., *BFI* = 1, we shall obtain *I*^*n*^(ROI_2_) = 0 and *I*^*n*^(ROI_1,3,4_) = 1 in the green channel, and *I*^*n*^(ROI_4_) = 0 and *I*^*n*^(ROI_1,2,3_) = 1 in the red channel. Therefore, a non-zero *BFI* value indicates that one or more streams were flowing back into the non-specific inlet channels and getting mixed with the other streams.

Figure 6b-d shows the normalized intensity plots for a relatively low total flow rate of 16.8 µl min^-1^, which corresponded to the most severe cases of backflow for Method A-C. At higher flow rates, the backflow effect was significantly mitigated (Figure S8). For Method A and *Q*_*t*_ of 16.8 µl min^-1^, the *I*^*n*^(ROI_2_) value was notably ≈ 0 for the green channel, whereas, the *I*^*n*^(ROI_4_) value for the red channel regularly dropped below 1 during the multiple stop flow cycles, leading to the highest *BFI* ≈ 0.28 recorded for any stop flow method (Figure 6b and 6e). At *Q*_*t*_ of 16.8 µl min^-1^, the *BFI* for Method A reduced to ≈ 0.16; however, it was still significantly higher than the other methods. Method B also struggled to stop the backflow with *BFI* ≈ 0.14 at *Q*_*t*_ of 16.8 µl min^-1^ as the *I*^*n*^(ROI_2_) and *I*^*n*^(ROI_4_) values for the green and red channels, respectively, regularly dropped below 1, whereas the remaining intensities increased above 0 over the three stop flow cycles recorded (Figure 6c). The *BFI* decreased to ≈ 0.06 as the flow rate was increased to *Q*_*t*_ of 168 µl min^-1^for Method B (Figure 6e). For Method C, the *I*^*n*^(ROI_2_) and *I*^*n*^(ROI_4_) values for the green and red channels, respectively, stayed close to 1 as desired, whereas the remaining intensities did not increase much above 0 over the three stop flow cycles recorded (Figure 6d). Comparatively, Method C performed better than the other two methods to restrict the backflow with *BFI* ≈ 0.04-0.05 over a broad range of flow rates (16.8-168 µl min^-1^).

## 3 Conclusions

We have successfully employed two-photon polymerization (2PP) printing to fabricate a 3D microfluidic device featuring a cascade of nozzles on a glass substrate. The 2PP printing enabled the fabrication of miniatured nozzles with a diameter of 75 µm, embedded within a main channel of 200 µm diameter. The microfluidic device converged four independently controlled flow streams into a single outlet after passing through a cascade of co-axial nozzles. As a result, a multi-layered co-axially sculpted cross-section flow profile was obtained. By carefully adjusting the flow rate ratio between individual streams, we could control the thicknesses of individual layers down to *O*(10µm) within the sculpted flow profile. We have investigated three different flow control methods, i.e., Method A-C, to rapidly stop the sculpted flow within the outlet of the microfluidic device. The best performing Method C could stop the flow by bringing the average flow velocity down by three orders of magnitude within ∼2s. Moreover, we observed during the stop-flow cycles that flow streams can leak backward into non-specific inlet channels due to the variable pressure drops from the inlets to the outlet and mismatched flow resistances. We defined a backflow index to quantify the backflow within the microfluidic device. We found that Method C performed relatively better compared to Methods A and B in terms of stopping the flow with the least delay time and minimizing the backflow with the smallest BFI values at variable total flow rates. Our findings highlight the complexity of stopping a microfluidic flow inside a multiple-inlet device with minimal backflow. Our findings will be of great benefit to realize efficient flow lithography processes for the fabrication of the next generation of functionalized materials.

## 4 Materials and methods

### 4.1 Device manufacturing

The substrate for the microfluidic chip consisted of 4-inch borosilicate glass wafers with a thickness of 1.1 mm (BOROFLOAT® 33 from Schott, Mainz, Germany), which are known for their excellent optical quality. A highly flexible laser micromachining system (microSTRUCT® C from 3D Micromac AG, Chemnitz, Germany) was used to locally roughen the glass surface, create through-holes and apply alignment marks for subsequent microchannel fabrication with a 2PP printer. The microSTRUCT® C system included a femtosecond (fs) laser (Pharos-10 W from Light Conversion Vilnius, Lithuania) emitting light at a fundamental wavelength of 1030 nm and operating at a pulse frequency of 100 kHz. Alignment marks were placed on both sides of the wafer. For each structuring task, the laser spot was guided by a galvanometer scanner (Scanlab RTC5, Puchheim, Germany) and moved along parallel lines at a speed of 1000 mm·s-^1^. After each complete pass, the fill structure was rotated 45°, and the laser focus was advanced 50 µm further into the substrate every fourth pass. The pulse energies used were 6.4 μJ for surface roughening and 22.6 μJ for drilling the through-holes (Table S6).

Microfluidic channels were fabricated on top of the roughened glass substrates using a 2PP system (Photonic Professional GT2, Nanoscribe GmbH, Germany), which allowed exposure of sub-micron voxels within a light-curing resist (Nanoscribe’s IPS negative tone photoresist, Young’s modulus: 5.1 GPa, refractive index: 1.515). The laser emitted pulses between 80-100 fs at a wavelength of 780 nm, with a repetition rate of 80 MHz. The 2pp system was configured for an average power of 50 mW at the standard objective aperture (LCI “Plan-Neofluar” 25×/0.8 Imm Korr Ph2 from Zeiss), with a power scaling factor of 1.0, allowing the laser power to be adjusted from 0% to over 100%. The microfluidic channel was printed at 45% power with a slicing distance (distance between each printed layer) of 1 μm and a hatching distance (distance between each voxel) of 0.5 µm, using a workspace of 300 × 300 × 300 μm^3^, with larger structures divided into blocks. A galvanometer scanner moved the laser laterally at 100 mm/s, and a piezo stage controlled the vertical movement between layers. The unpolymerised photoresist was removed using an mr-Dev 600 developer bath, and any residual within the channels was extracted by suction. After drying, the printed structures were cured on a hotplate at 190°C for 10 minutes, resulting in high transparency, hydrophobic surfaces, and strong adhesion due to the locally roughened glass substrate (Table S7).

### 4.2 Experimental set-up

The microfluidic device was connected to the syringe pumps (Nemesys S, CETONI Gmbh) by using a self-designed two-part device holder with adequate ports for fluidic connections at the bottom. The device holder was composed of a micromachined aluminum base and lid, which were screwed (M3) together to sandwich the microfluidic device. The inlet and outlet ports within the base of the holder connected the microfluidic device to the fluid supply and waste tube by using male connectors (M6, TechLab) and PTFE tubing (1/16” – 1 mm OD-ID, TechLab). The fluid flow from the syringes to the device inlets and form device outlet to the waste collection tube (15 mL PP tube, CELLSTAR®, Greiner Bio-One International GmbH) was controlled by 3-way valves (ASCO 833-630887, 3 bar, EMERSON Electric Co.). The syringe pumps and the valves were connected to an I/O-B module (Modular Qmix I/O-B Module, Cetoni) and controlled using a custom LabVIEW (National Instrument) script for the cyclic operation of stop-flow methods.

Matching flow resistances: Additional flow resistances were added to the reservoir side by using PTFE tubings (1/16” mm OD, TechLab) with 250 µm inner diameter. The length was adjusted according to the pressure drop inside the microfluidic inlets. For inlets 1-3, the additional tubing length was 60 cm, whereas for inlet 4, the tubing length was 90cm. This resulted in the added hydraulic flow resistances as *R*_*1-3*_ = 2.19 × 10^14^ Pa.s/m^3^ and *R*_*4*_ = 3.29 × 10^14^ Pa.s/m^3^.

Sample preparation: Poly(ethylene glycol) diacrylate (PEGDA, Mn:575, Sigma–Aldrich) was diluted with deionized (DI) water at a 60% volume ratio and sterile filtered (1.0 µm, Target2™, Thermo Scientific) for all experiments. The solution was mixed with fluorescent tracing particles (PS-FluoGreen-5.0, Microparticles Gmbh) at ≈ 10^6^ particles per milliliter.

Confocal microscopy: Confocal imaging was conducted with a concentration of 0.2 mmol/L Rhodamine B and with a concentration of 0.32 mmol/ L Fluorescein isothiocyanate separately mixed with a 60% PEGDA- EtOH solution.

### 4.3 Backflow quantification by particle velocity measurement

Polystyrene green fluorescent particles (PS-FluoGreen-5.0, 5µm, Microparticles Gmbh) were suspended in the liquid sample for the flow velocity measurement inside the microchannel. The central region of the microchannels (region of interest (ROI): 200 µm channel width x 2600 µm length) was observed by using a fluorescent microscope (Thunder Imager DMi8, Leica Microsystems) equipped with an integrated camera (DFC9000, Leica Microsystems) capturing an image every 40 ms (i.e., 25 fps) through a 5x magnification objective. Following a waiting period of at least four stop-flow cycles to pressurize the system, the particles were recorded over three consecutive cycles. A custom-written computer code (MATLAB R2021a, The MathWorks Inc.) was employed to extract the particle velocities from the recorded movies. The particles were segmented based on gray value thresholding, and their centroids were calculated. Subsequently, the distances (*s*_p_) and angles (*α*_p_) between each particle from frame f_i_ and consecutive frame f_i+1_ were surjectively calculated, whereby the angle was determined with respect to a vector parallel to the channel walls. Matching particles were determined by minimizing a loss function, considering normal and backflow while penalizing large distances and off-streamline angles:

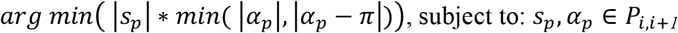

where *P*_1,1+*1*_ denotes the domain of particle pairs in consecutive frames. The individual particle’s velocity vector was obtained by multiplying the distance with the frame rate. Due to the parabolic flow profile, particles located near the microchannel walls were excluded. Once the individual particle velocities were determined for each cycle, the velocity data were overlaid for the three cycles to calculate an average residual flow velocity.

## 5 Acknowledgments

This work was supported by German Research Foundation (DFG, project # 494539689).

## 7 Supplementary information

**Figure S1:**
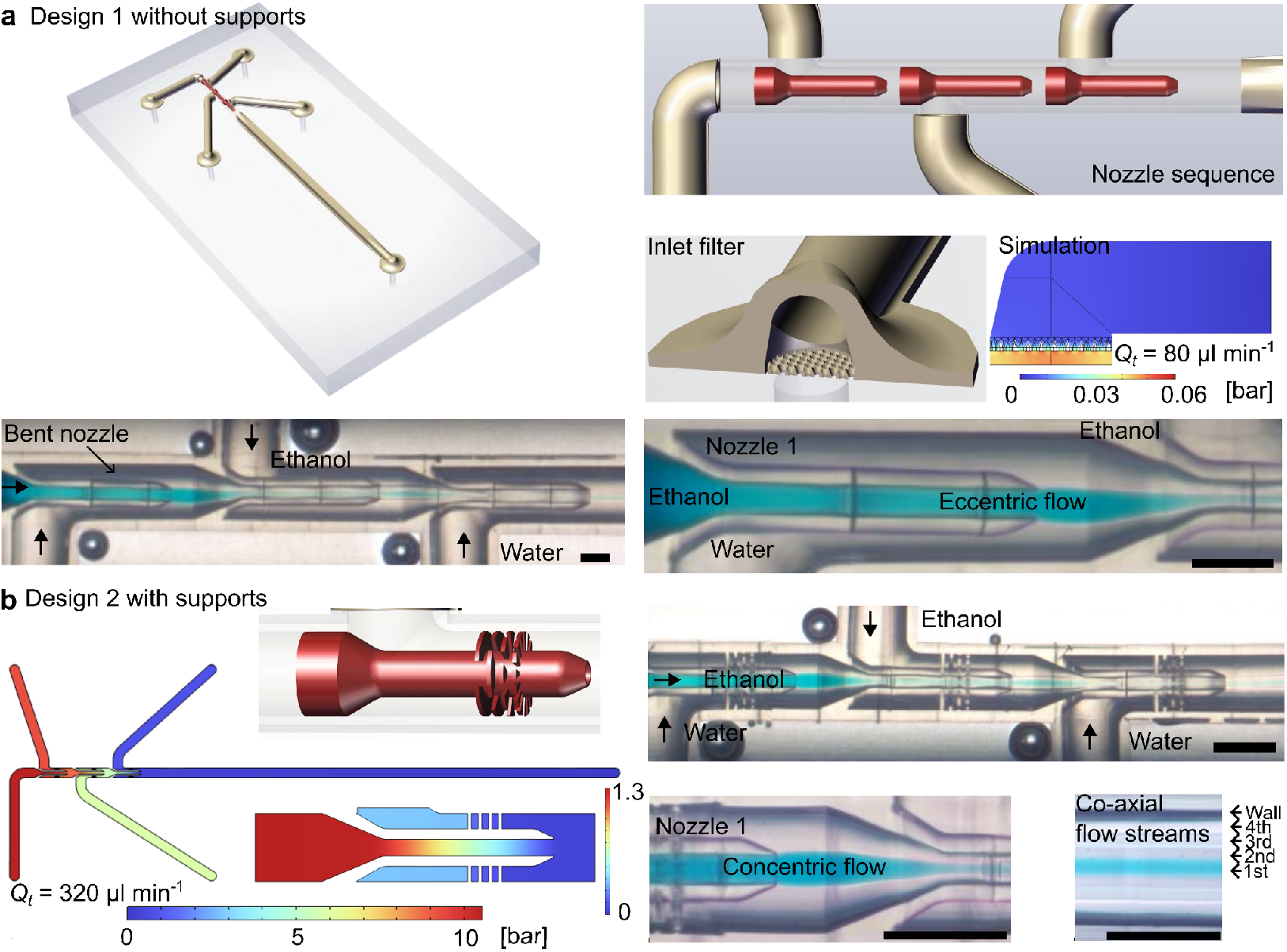
Device development. (a) Preliminary device design 1 without support structure for the nozzles. The free-standing nozzles bent toward the sidewalls over time due to the inhomogeneous fluid pressure around the nozzle. The bent nozzles produced an eccentric flow of co-flowing water and ethanol streams. The innermost stream was mixed with a blue dye to trace the flow. An inlet filter, introduced to restrict the flow of undesired particles into the channel, added to the flow resistance and pressure drop in the fluidic circuit. (b) In a revised version of the device, support structures were added to stabilize the nozzles, which resulted in a concentric coaxial flow of four streams (ethanol-water-ethanol-water). However, the added nozzle supports contributed to a high fluidic resistance and undesirable pressure drop over the nozzle length. For a viscous polymer sample (10x more viscous than water), a pressure-drop of up to 10 bar was estimated using the numerical simulation due to the smaller inner diameter of the nozzles, i.e., 50 μm.

**Figure S2:**
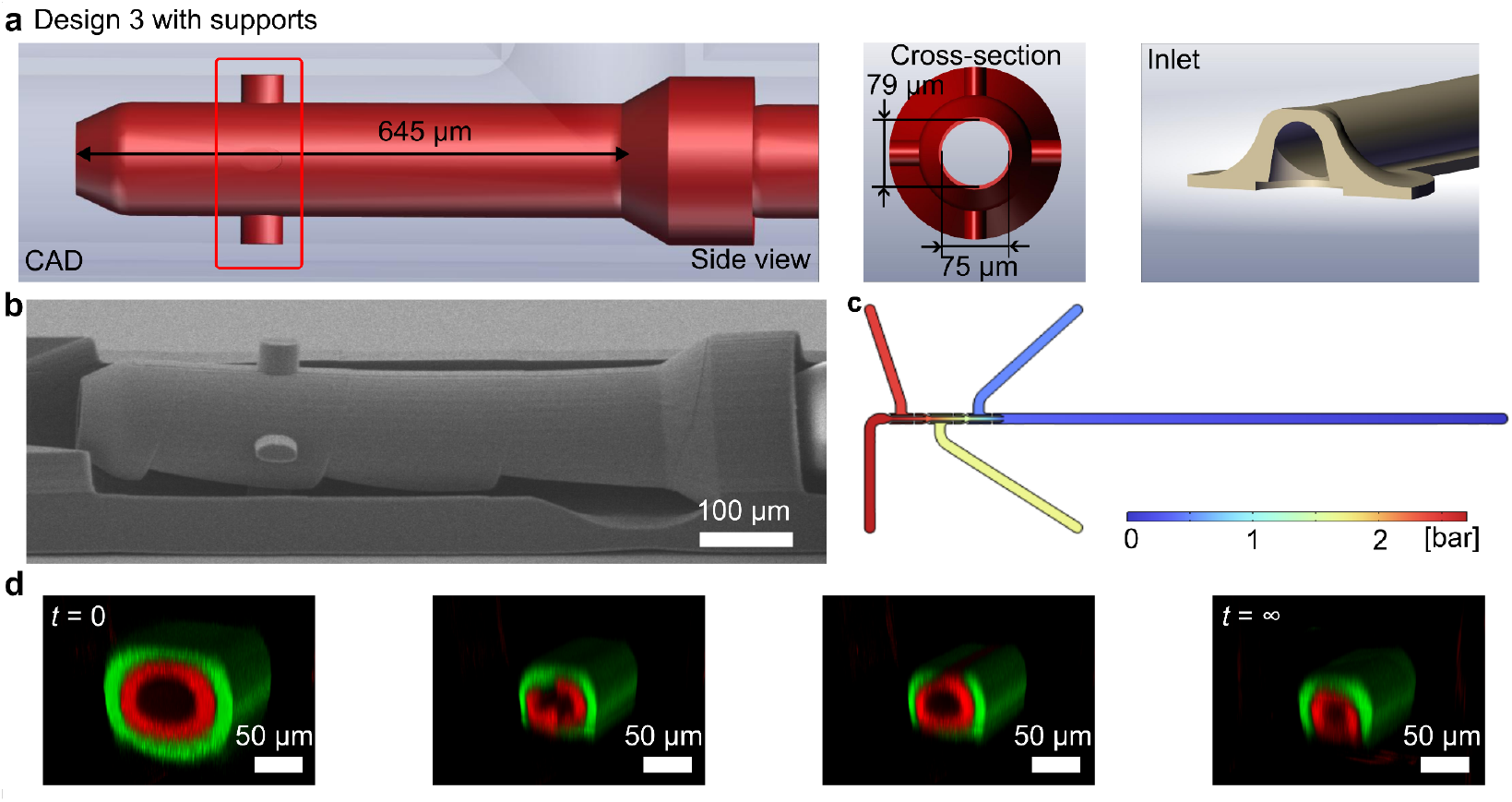
Microfluidic channel design 3. (a) A single cross-bar support structure to keep the pressure drop at low as possible. The nozzle’s cross-sectional dimensions were adjusted to compensate for the ellipsoidal voxel shape of the 3D print. The inlet filers were also removed. (B) SEM images showed the nozzle sagging at multiple locations in the absence of multiple adequate support structure. (C) Confocal imaging of sculpted co-axial flow inside the microfluidic device traced by two fluorescently labeled (green and red) streams sandwiched by two transparent streams. Flow structure changed over time due to the unstable and under-supported nozzles.

**Figure S3:**
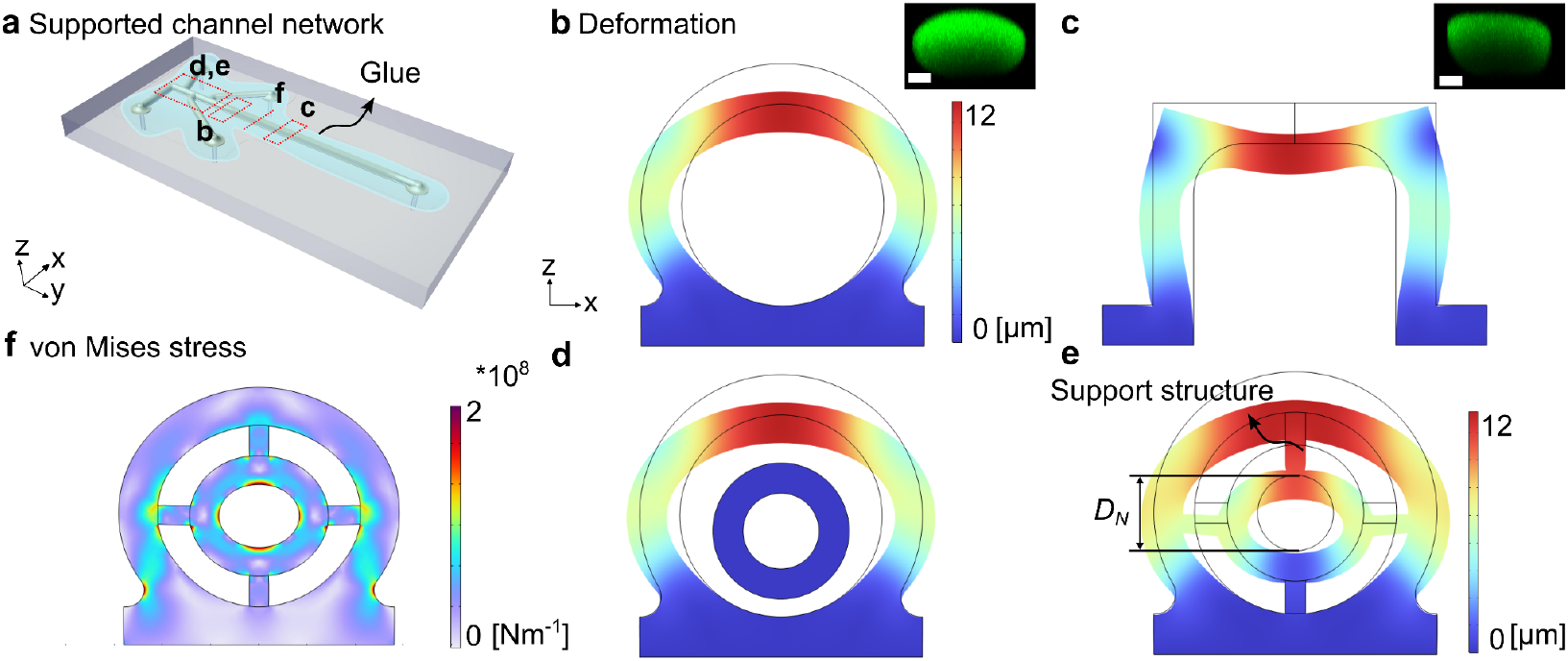
Microfluidic channel and nozzle deformation. (a) A layer of protective glue was added at the top of the microfluidic channel network to give additional support to to 3D printed structures. (b-c) Numerical simulations showed the channel deformation under an applied load (2.5^*^10^7^ Nm^-1^) in the negative z-direction, which mimicked the stress applied by the UV-cured protective resin. In the insets, the confocal images of the microchannel, filled with a green fluorescent dye, at two different locations showed a clear deviation from the originally designed circular and square channel cross-sections. (d-f) The support structures assisting the nozzle were also contributing to the change in its cross-sectional shape as the outer channel wall was deformed under an applied load. As a comparison, the nozzle without the support structure was not deformed (d) even when the outer channel was deformed.

**Figure S4:**
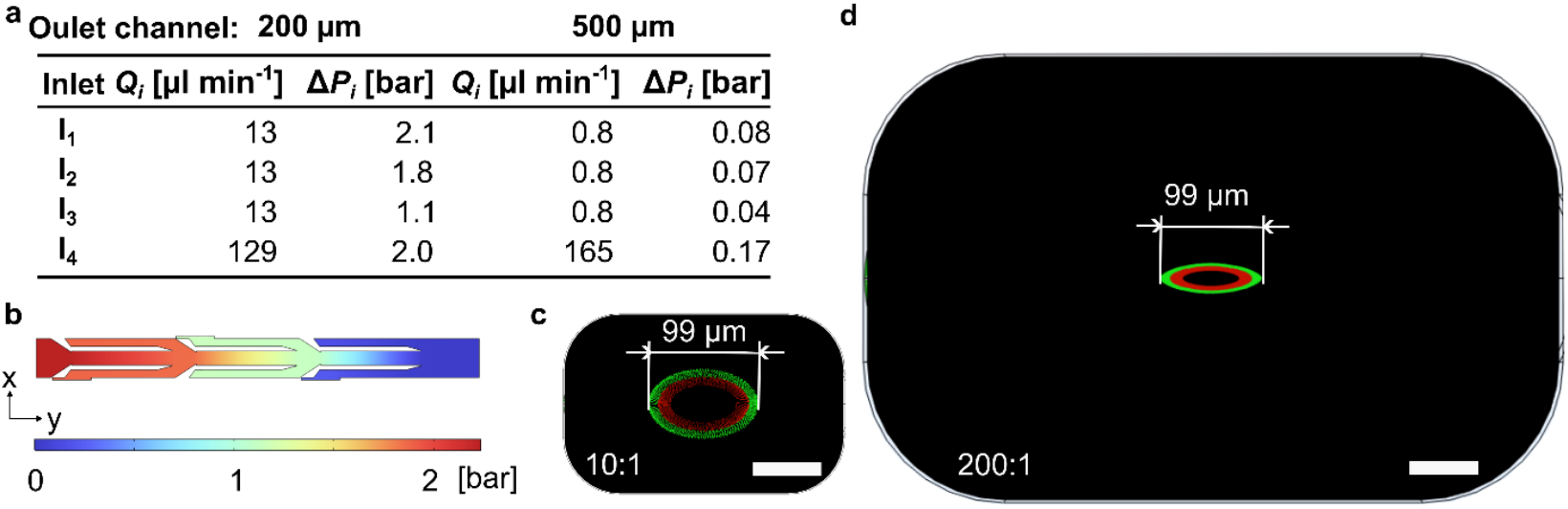
(a) Numerical simulations of multilayered flow inside microchannels with nominal hydraulic diameter (*D*_*h,c*_) of ∼200 µm amd ∼500 µm. The corresponding flow rates and the pressure drop values for the four inlets are listed. (b) Pressure contour for the device with *D*_*h,c*_ ≈ 200 µm. (c) Inside the miniaturized device with *D*_*h,c*_ ≈ 200 µm, a flow rate ratio of *Q*_*4*_:*Q*_*1-3*_ = 10:1 resulted in a diameter of the green layer *D*_*g*_ = 99 µm. (d) Inside the 2.5x larger device with *D*_*h,c*_ ≈ 500 µm, a flow rate ratio of *Q*_*4*_:*Q*_*1-3*_ = 200:1 resulted in a diameter of the green layer *D*_*g*_= 99 µm. The scale bars are 50 µm.

**Figure S5:**
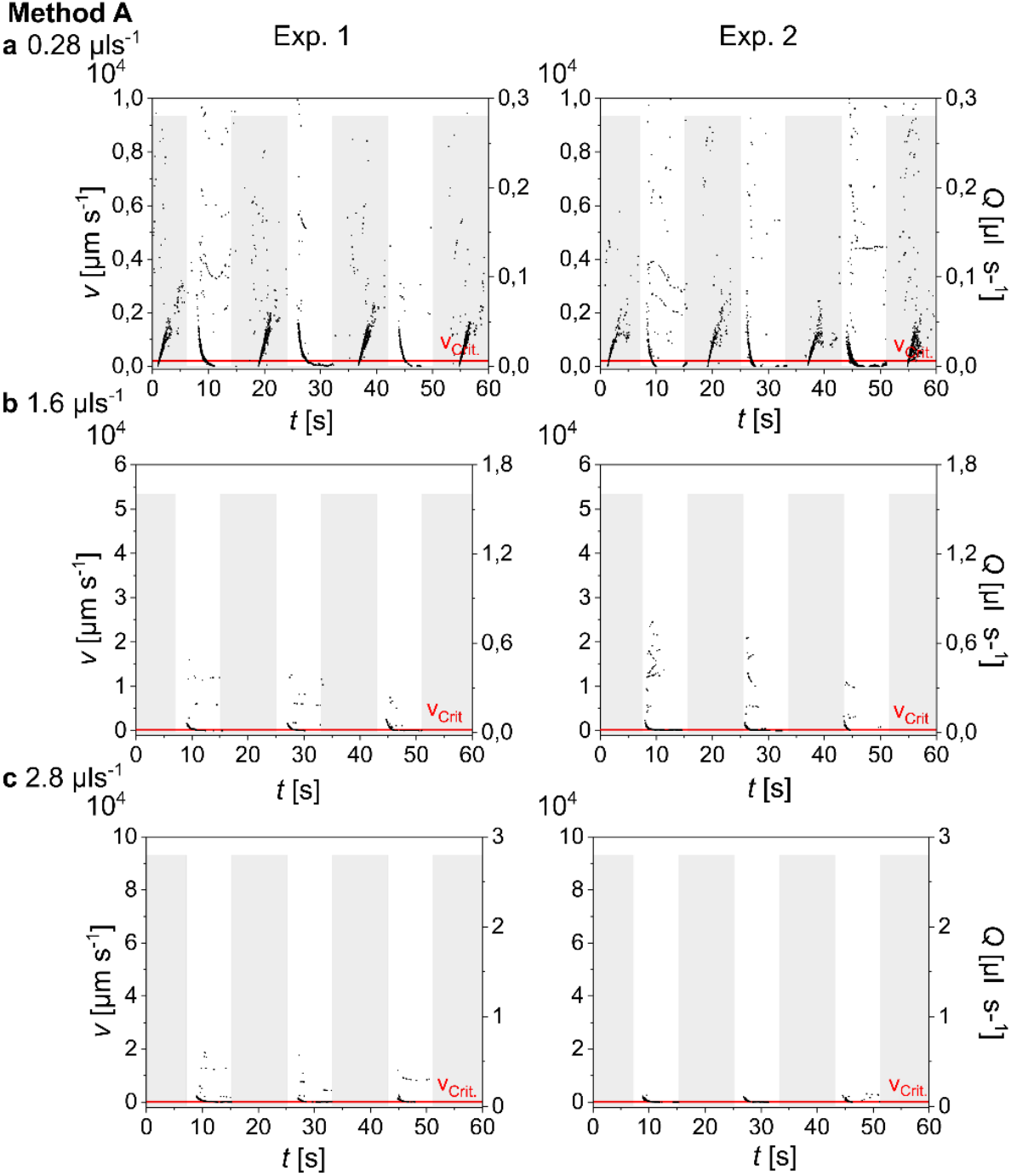
Stop-flow residual velocity plots for Method A for three different flow rates (0.28, 1.6, 2.8 µl s^-1^) or (16.8, 96, 168 µl min^-1^). It was noted that the higher flow rates could stabilize the flow relativley quickly within the microchannel compared to lower flow rates.

**Figure S6:**
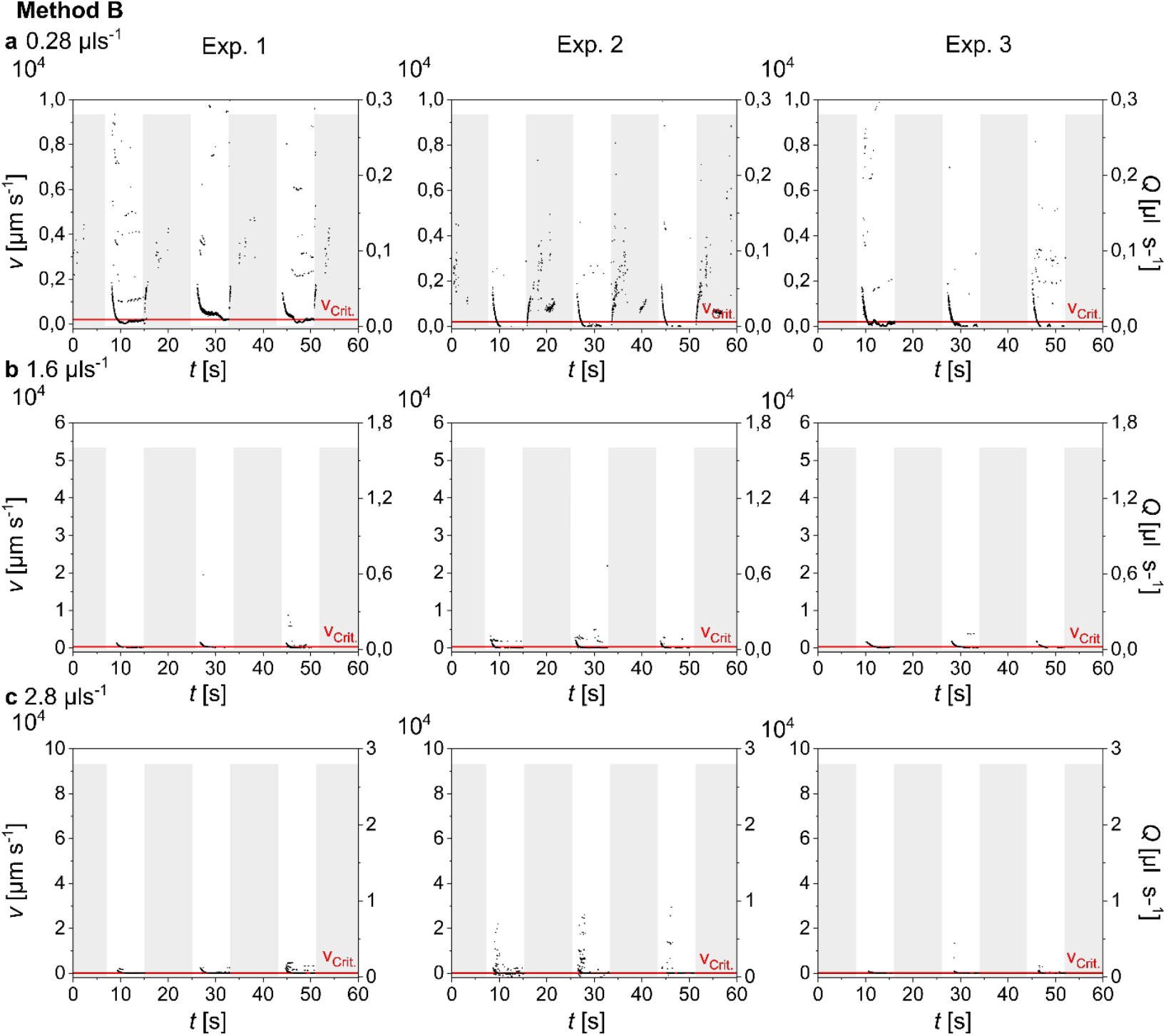
Stop-flow residual velocity plots for Method B for three different flow rates (0.28, 1.6, 2.8 µl s^-1^) or (16.8, 96, 168 µl min^-1^).

**Figure S7:**
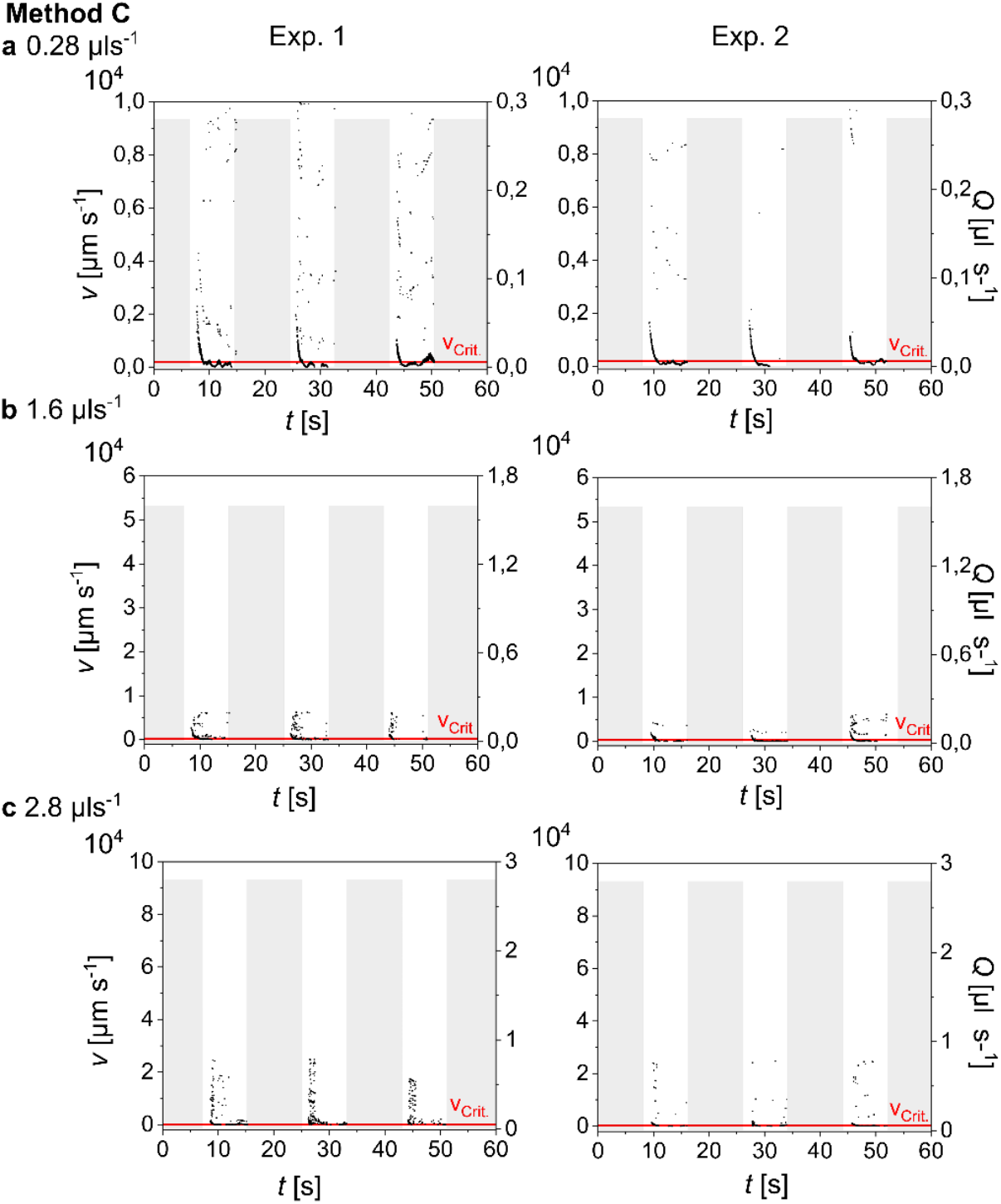
Stop-flow residual velocity plots for Method C for three different flow rates (0.28, 1.6, 2.8 µl s^-1^) or (16.8, 96, 168 µl min^-1^).

**Figure S8:**
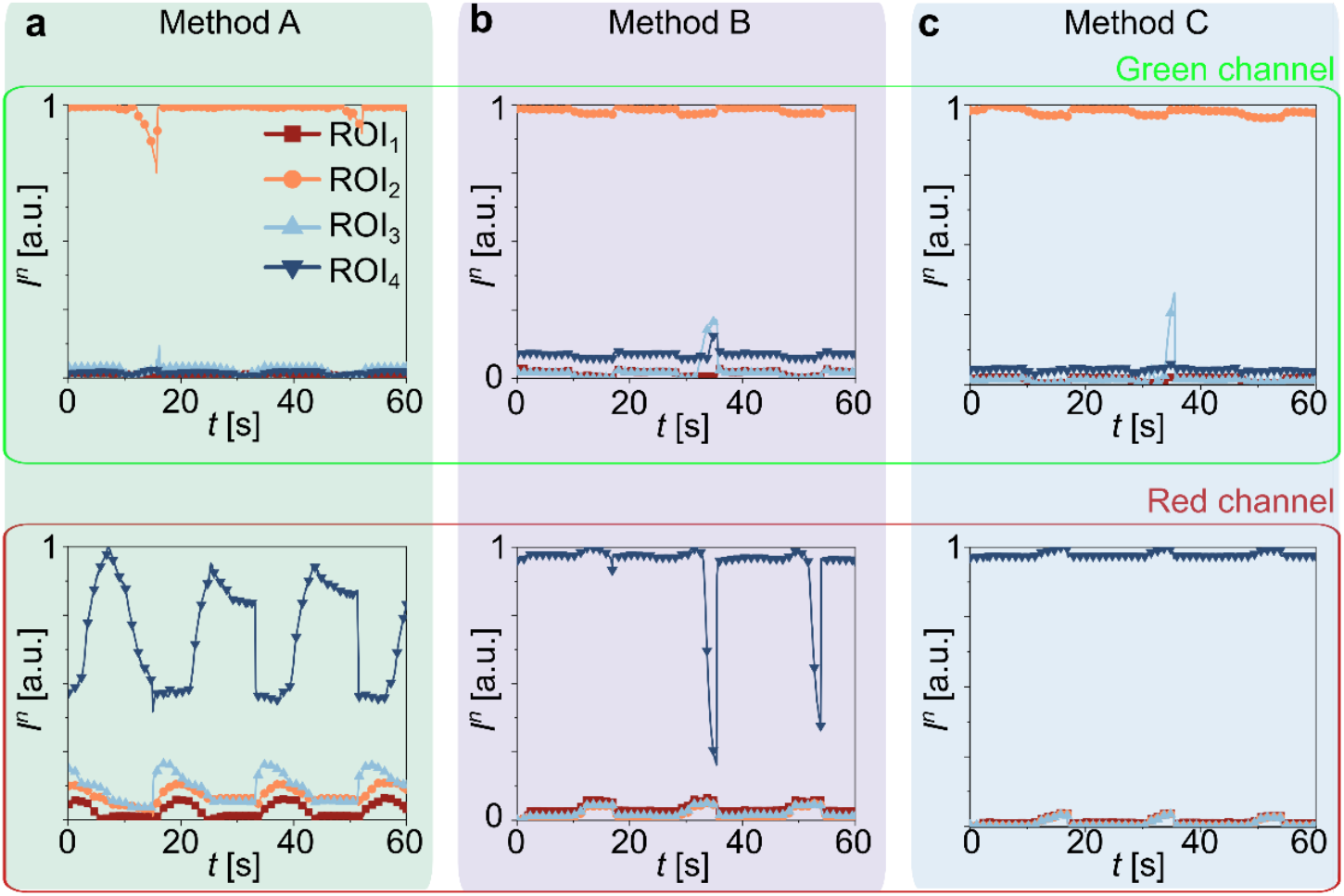
Backflow inside the microfluidic device during the stop-flow cycle. (a-c) Fluorescence intensity at four different ROIs for Methods A-C with a total flow rate of 2.8 µl s^-1^ or 168 µl min^-1^.

**Figure S9:**
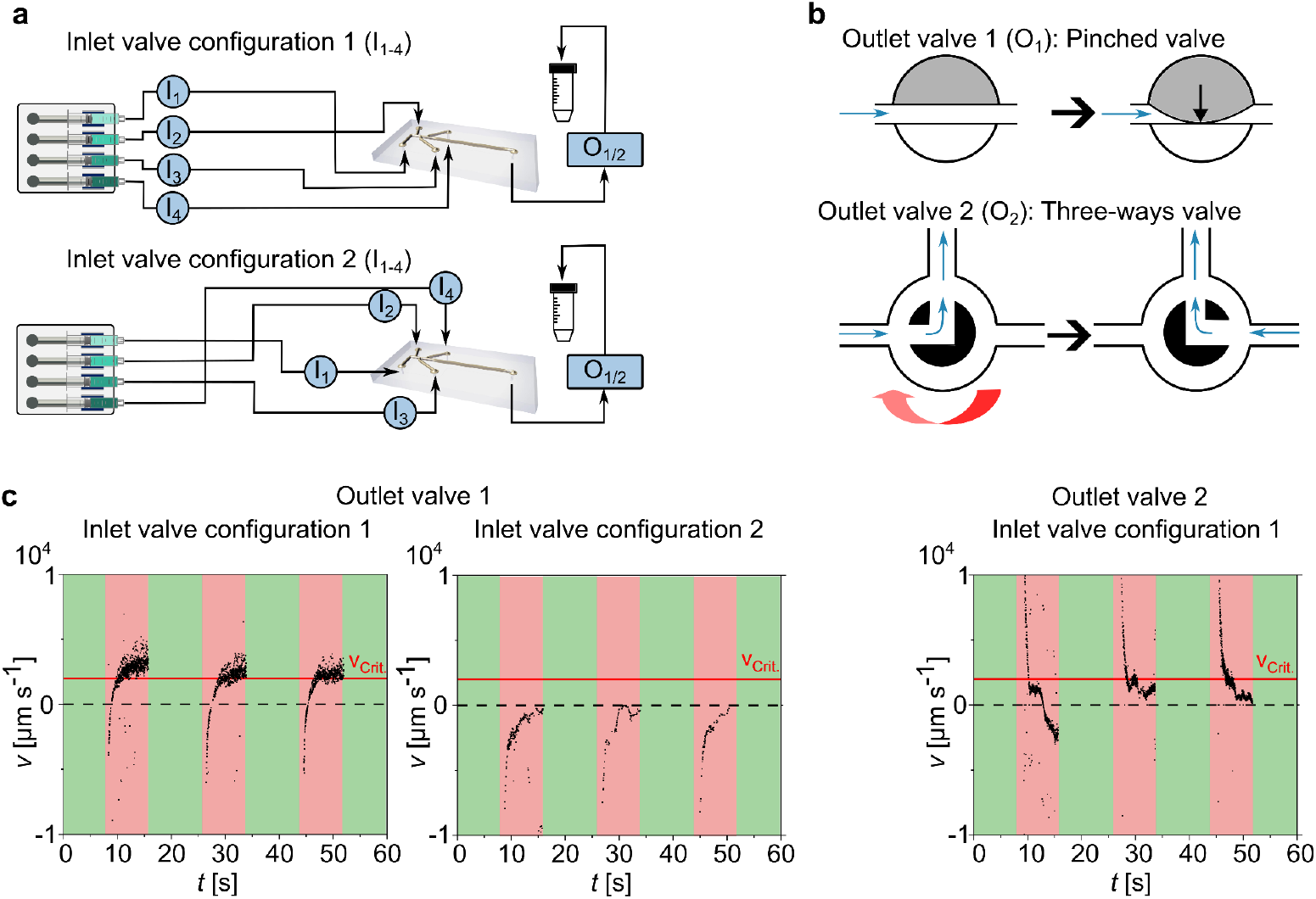
Investigation of different inlet valve positions and outlet valve types. (a) Different inlet valve positions, either in front of the microchannel or in front of the syringes. (b) Principle of both outlet valve types, i.e., pinched valve and three-way valve. (C) Residual flow velocity over time from three stop-flow cycles with different combinations. Outlet valve 1 (pinched valve) combined with inlet valves in front of the syringe (inlet valve configuration 1) or in front of the microchannel (inlet valve configuration 2). The pinched valve was not suitable to stop the flow in a reasonable time because of backflow during the stop phase. The valve in front of the syringe was easier to adapt in the circuit due to the limited space on the microscope table. Outlet valve 2 (three-way valve) and inlet valve configuration 1 were the most suitable for further stop-flow investigations.

## Movies captions

**Movie S1**: Stopping multi-layered co-axial flow inside the microchannel for three different stop-flow methods A-C for three different total flow rates of 16.8, 96, 168 µl min^-1^.

**Movie S2**: Backflow analysis using green and red fluorescent dyed streams inside the microchannel for three different stop-flow methods A-C.

**Table S1:** Important experimental parameters and results of pressure drop simulations with microfluidic device design 2, which had supports and inlet filters.

| $Q_t$<br>[$\mu\text{l min}^{-1}$] | Ratio $Q_4:Q_{1-3}$ | $Re$ | $\eta$ [Pa s] | $D_{h,c}$<br>[mm] | $D_{h,n}$<br>[mm] | $L_n$ [mm] | $\Delta P_{filter}$<br>[bar] | $\Delta P_{tubing}$<br>[bar] | $\Delta P_{noz. in}$<br>[bar] | $\Delta P_{noz. out}$<br>[bar] | $\Delta P_{1,2,3,4}$<br>[bar] | $\Delta P_{max}$<br>[bar] | $Pe$ | $D$ [$\text{m}^2 \text{s}^{-1}$] |
| --- | --- | --- | --- | --- | --- | --- | --- | --- | --- | --- | --- | --- | --- | --- |
| 16.8 | 2:1 | 0.05 | 0.035 | 0.2 | 0.05 | 0.5 | 0.003 | $3.0 \cdot 10^{-3}$ | 0.07 | 0.02 | 0.06, 0.25,<br>0.39, 0.45 | **0.45** | $1.78 \cdot 10^4$ | $10^{-10}$ |
| 40 | 2:1 | 0.12 | 0.035 | 0.2 | 0.05 | 0.5 | 0.006 | $7.2 \cdot 10^{-3}$ | 0.17 | 0.04 | 0.13, 0.60,<br>0.93, 1.06 | **1.07** | $4.24 \cdot 10^4$ | $10^{-10}$ |
| 320 | 2:1 | 0.96 | 0.035 | 0.2 | 0.05 | 0.5 | 0.048 | $57.0 \cdot 10^{-3}$ | 1.33 | 0.33 | 1.07, 4.78,<br>7.48, 8.50 | **8.55** | $3.40 \cdot 10^5$ | $10^{-10}$ |

**Table S2:** Important experimental parameters and results of pressure drop simulations with microfluidic device design 3, which had supports but no inlet filters.

| $Q_t$<br>[$\mu\text{l min}^{-1}$] | Ratio $Q_4:Q_{1-3}$ | $Re$ | $\eta$ [Pa s] | $D_{h,c}$ [mm] | $D_{h,n}$<br>[mm] | $L_n$<br>[mm] | $\Delta P_{filter}$<br>[bar] | $\Delta P_{tubing}$<br>[bar] | $\Delta P_{noz. in}$<br>[bar] | $\Delta P_{noz. out}$<br>[bar] | $\Delta P_{1,2,3,4}$<br>[bar] | $\Delta P_{max}$<br>[bar] | $Pe$ | $D$ [$\text{m}^2 \text{s}^{-1}$] |
| --- | --- | --- | --- | --- | --- | --- | --- | --- | --- | --- | --- | --- | --- | --- |
| 40 | 2:1 | 0.12 | 0.035 | 0.2 | 0.075 | **0.675** | 0 | $7.2 \cdot 10^{-3}$ | 0.04 | 0.01 | 0.08, 0.18,<br>0.25, 0.28 | **0.28** | $4.24 \cdot 10^4$ | $10^{-10}$ |
| 80 | 2:1 | 0.24 | 0.035 | 0.2 | 0.075 | **0.675** | 0 | $14.2 \cdot 10^{-3}$ | 0.08 | 0.03 | 0.15, 0.36,<br>0.51, 0.56 | **0.56** | $8.49 \cdot 10^4$ | $10^{-10}$ |
| 160 | 2:1 | 0.48 | 0.035 | 0.2 | 0.075 | **0.675** | 0 | $28.6 \cdot 10^{-3}$ | 0.15 | 0.05 | 0.31, 0.72,<br>1.02, 1.11 | **1.11** | $1.70 \cdot 10^5$ | $10^{-10}$ |
| 320 | 2:1 | 0.96 | 0.035 | 0.2 | 0.075 | **0.675** | 0 | $57.0 \cdot 10^{-3}$ | 0.31 | 0.11 | 0.62, 1.43,<br>2.04, 2.23 | **2.24** | $3.40 \cdot 10^5$ | $10^{-10}$ |
| 168 | 1:1 | 0.50 | 0.035 | 0.2 | 0.075 | **0.675** | 0 | $29.9 \cdot 10^{-3}$ | 0.20 | 0.07 | 1.42, 1.29,<br>0.90, 0.27 | **1.42** | $1.78 \cdot 10^5$ | $10^{-10}$ |

\*Updated and calculated using online pressure drop calculator (https://www.pressure-drop.com/Online-Calculator/) for a tubing with 75 cm length and 1 mm diameter density 987 kg/m<sup>3</sup> and viscosity 35 mPa

**Table S3:** Horizontal dimensions of the sculpted flow profile depending on the flow rate ratio for both support structures.

| Support structure 1 |  |  |  |  |  | Support structure 2 |  |  |  |
| --- | --- | --- | --- | --- | --- | --- | --- | --- | --- |
| $Q_4:Q_3:Q_2:Q_1$ | $Q_t$ [$\mu\text{l min}^{-1}$] | $D_o$ [$\mu\text{m}$] | $D_c$ [$\mu\text{m}$] | $t_{out}$ [$\mu\text{m}$] | $t_{in}$ [$\mu\text{m}$] | $D_o$ [$\mu\text{m}$] | $D_c$ [$\mu\text{m}$] | $t_{out}$ [$\mu\text{m}$] | $t_{in}$ [$\mu\text{m}$] |
| 1:1:1:1 | 168 | 150 | 48 | 17 | 23 | 141 | 59 | 20 | 29 |
| 2:1:1:1 | 168 | 131 | 43 | 19 | 19 | 124 | 45 | 18 | 31 |
| 10:1:1:1 | 168 | 80 | 30 | 9 | 11 | 71 | 33 | 12 | 14 |
| 1:2:1:1 | 168 | 156 | 56 | 30 | 21 | 149 | 67 | 32 | 18 |
| 1:1:2:1 | 168 | 155 | 33 | 14 | 30 | 152 | 36 | 17 | 41 |
| 2:1:1:2 | 168 | 136 | 72 | 12 | 12 | 131 | 58 | 13 | 17 |
| 1:2:2:1 | 168 | 160 | 25 | 24 | 29 | 154 | 39 | 27 | 34 |

**Table S4:** Horizontal dimensions of the sculpted flow profile depending on the flow rate ratio for both support structures.

| Support structure 1 |  |  |  |  |  | Support structure 2 |  |  |  |
| --- | --- | --- | --- | --- | --- | --- | --- | --- | --- |
| $Q_4:Q_3:Q_2:Q_1$ | $Q_t$ [$\mu\text{l min}^{-1}$] | $D_o$ [$\mu\text{m}$] | $D_c$ [$\mu\text{m}$] | $t_{out}$ [$\mu\text{m}$] | $t_{in}$ [$\mu\text{m}$] | $D_o$ [$\mu\text{m}$] | $D_c$ [$\mu\text{m}$] | $t_{out}$ [$\mu\text{m}$] | $t_{in}$ [$\mu\text{m}$] |
| 1:1:1:1 | 168 | 123 | 37 | 22 | 12 | 118 | 30 | 18 | 25 |
| 2:1:1:1 | 168 | 105 | 31 | 25 | 16 | 102 | 23 | 20 | 13 |
| 10:1:1:1 | 168 | 59 | 15 | 0 | 13 | 69 | 14 | 8 | 9 |
| 1:2:1:1 | 168 | 135 | 50 | 36 | 16 | 130 | 24 | 30 | 23 |
| 1:1:2:1 | 168 | 122 | 35 | 27 | 17 | 115 | 14 | 14 | 41 |
| 2:1:1:2 | 168 | 97 | 30 | 30 | 21 | 105 | 31 | 17 | 20 |
| 1:2:2:1 | 168 | 131 | 28 | 20 | 16 | 126 | 14 | 32 | 26 |

**Table S5:** Deviation in the sculpted flow profile dimensions.

| Support structure 1 |  |  |  |  |  | Support structure 2 |  |  |  |
| --- | --- | --- | --- | --- | --- | --- | --- | --- | --- |
| $Q_4:Q_3:Q_2:Q_1$ | $Q_t$ [$\mu\text{l min}^{-1}$] | $\Delta D_o$ [%] | $\Delta D_c$ [%] | $\Delta t_{out}$ [%] | $\Delta t_{in}$ [%] | $\Delta D_o$ [%] | $\Delta D_c$ [%] | $\Delta t_{out}$ [%] | $\Delta t_{in}$ [%] |
| 1:1:1:1 | 168 | 10 | 13 | -13 | 32 | 9 | 33 | 5 | 6 |
| 2:1:1:1 | 168 | 11 | 16 | -13 | 8 | 9 | 34 | -5 | 41 |
| 10:1:1:1 | 168 | 15 | 33 | 1 | -7 | 2 | 40 | 23 | 25 |
| 1:2:1:1 | 168 | 7 | 6 | -9 | 14 | 7 | 48 | 4 | -11 |
| 1:1:2:1 | 168 | 12 | 3 | -32 | 29 | 14 | 43 | 11 | 0 |
| 2:1:1:2 | 168 | 17 | 42 | -42 | -27 | 11 | 30 | -12 | -8 |
| 1:2:2:1 | 168 | 10 | 5 | 9 | 29 | 10 | 47 | -9 | 14 |

**Table S6:** Parameters for glass substrate treatment.

| Parameter (fs laser ) | Through Hole Creation | Roughening |
| --- | --- | --- |
| Wavelength | 1030 nm | 1030 nm |
| Pulse Frequency | 100 kHz | 100 kHz |
| Pulse Energy | 6.4 μJ | 22.6 μJ |
| Laser Spot Movement Speed | 1000 mm/s | 1000 mm/s |
| Scanning Strategy | Parallel lines, rotate 45° after one pass | Parallel lines, rotate 45° after one pass |
| Focus Adjustment | Move 50 μm deeper every fourth pass | |

**Table S7:**
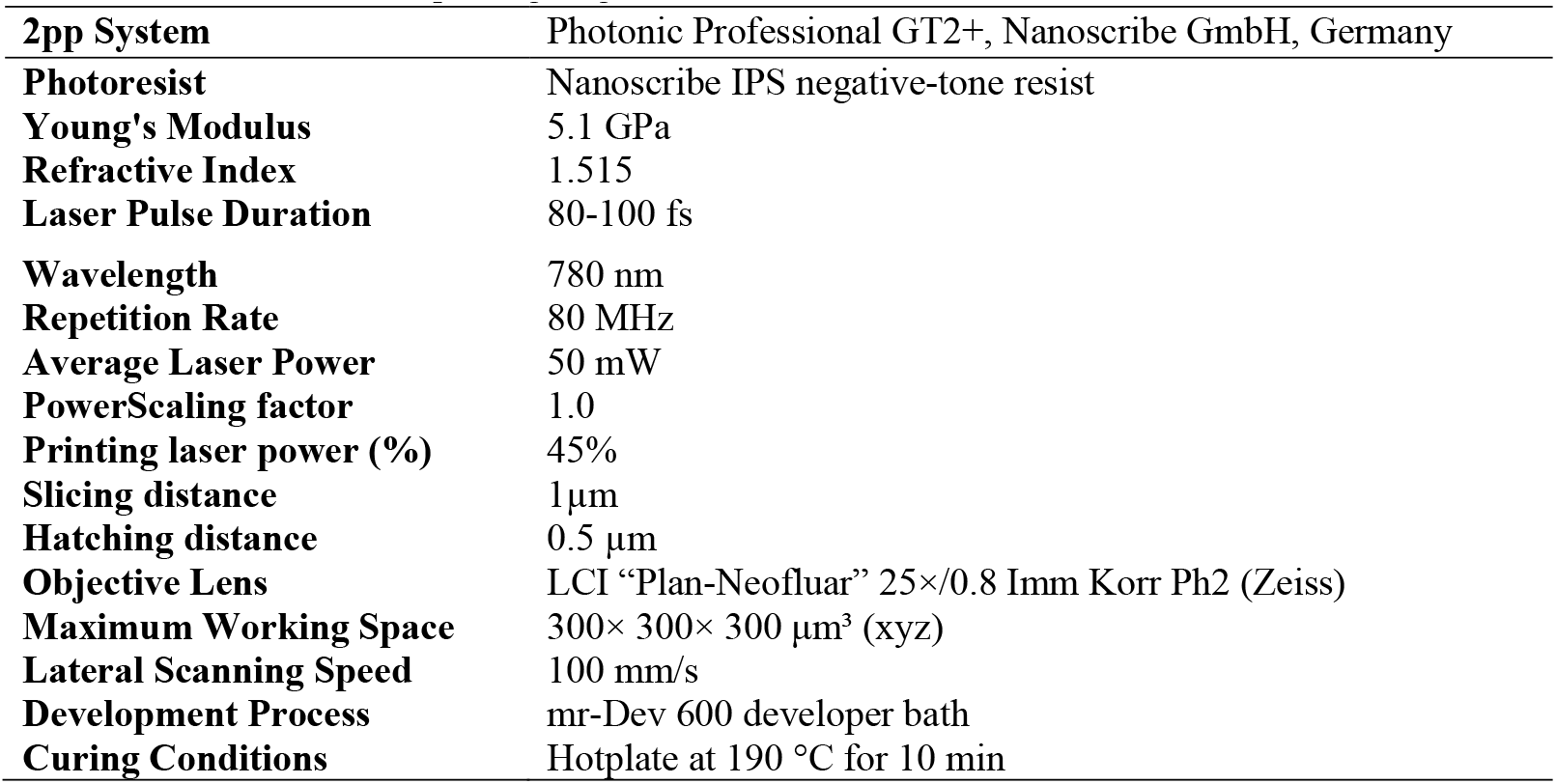
Parameters for 2PP printing on glass substrate.

